# Cancer causes dysfunctional insulin signaling and glucose transport in a muscle-type specific manner

**DOI:** 10.1101/2021.11.03.467058

**Authors:** Steffen H. Raun, Jonas Roland Knudsen, Xiuqing Han, Thomas E. Jensen, Lykke Sylow

## Abstract

Metabolic dysfunction and insulin resistance are emerging as hallmarks of cancer and cachexia, and impair cancer prognosis. Yet, the molecular mechanisms underlying impaired metabolic regulation is not fully understood. To elucidate the mechanisms behind cancer-induced insulin resistance in muscle, we isolated *extensor digitorum longus* (EDL) and soleus muscles from Lewis Lung Carcinoma tumor-bearing mice. Three weeks after tumor inoculation, muscles were isolated and stimulated with or without a submaximal dose of insulin (1.5 nM). Glucose transport was measured using 2-[^3^H]Deoxy-Glucose and intramyocellular signaling was investigated using immunoblotting. In soleus muscles from tumor-bearing mice, insulin-stimulated glucose transport was abrogated concomitantly with abolished insulin-induced TBC1D4 and GSK3 phosphorylation. In EDL, glucose transport and TBC1D4 phosphorylation were not impaired in muscles from tumor-bearing mice, while AMPK signaling was elevated. Anabolic insulin signaling via phosphorylation of the mTORC1 targets, p70S6K thr389 and ribosomal-S6 ser235, were decreased by cancer in soleus muscle while increased or unaffected in EDL. In contrast, the mTOR substrate, pULK1 ser757, was reduced in both soleus and EDL by cancer. Hence, cancer causes considerable changes in skeletal muscle insulin signaling that is dependent of muscle-type, which could contribute to metabolic dysregulation in cancer. Thus, skeletal muscle could be a target for managing metabolism in cancer.

**Highlights:**

- Cancer abrogates insulin-stimulated glucose transport selectively in oxidative soleus muscle
- Multiple TBC1D4 phosphorylation sites are reduced in cancer-associated muscle insulin resistance
- Cancer leads to increased AMPK signaling in the glycolytic EDL muscle
- Cancer alters anabolic insulin signaling in soleus and EDL muscle

## 1.0 Introduction

Within the last decades, it has become evident that cancer causes severe systemic alterations of the host. While unwanted loss of skeletal muscle and fat mass, coined cachexia^1^ is well-described, a lesser described burden of many cancers is the severe metabolic dysregulation. Evidently, several cancers, and in particular cachexia-inducing cancers, are associated with poor metabolic regulation, including insulin resistance in both pre-clinical models^2–4^ and human patients^5–9^. While the underlying mechanisms are still poorly defined, they are crucial to delineate, as dysregulated metabolism is highly associated with cancer incidence, poor cancer prognosis, and increased recurrence rates^10–15^.

Skeletal muscle insulin resistance and dysregulated metabolism are detrimental to whole body glucose homeostasis, as skeletal muscle is responsible for the majority of insulin-stimulated glucose disposal^16^. We recently showed, that cancer causes severe insulin resistance in pre-cachectic tumor-bearing mice^3^ on several parameters, including reduced skeletal muscle and white adipose tissue glucose uptake and abrogated insulin-stimulated microvascular perfusion^3^. Yet, the muscle-specific contributions and molecular defects were not identified in that study. In addition, it is unknown whether different muscle-types, Type I fiber- or Type II fiber-dominated muscles, are affected by cancer to a similar degree with regards to insulin resistance towards glucose uptake and anabolism.

To elucidate the muscle-intrinsic mechanisms that contribute to skeletal muscle insulin resistance in cancer, we here conducted a detailed investigation of glucose uptake and intramyocellular signaling in response to insulin in isolated oxidative (Type I fiber-dominated) and glycolytic (Type II fiber-dominated) muscles from tumor-bearing mice. It was hypothesized that muscles isolated from tumor-bearing mice would display altered insulin signaling leading to decreased glucose transport and anabolism.

## 2.0 Materials and Methods

### 2.1 Animals and ethics

A total of 28 C57bl6/J (Taconic, Lille Skensved, DK) mice, 12 weeks old, female, were group housed at ambient temperature (21-23 °C) with nesting materials. The mice were held on a 12 h:12 h light-dark cycle with access to a standard rodent chow diet (Altromin no. 1324, Brogaarden, DK) and water *ad libitum*. All experiments were approved by the Danish Animal Experimental Inspectorate (Licence: 2016-15-0201-01043). The sample size (n=>10 in each condition) were decided from previous work with the experimental incubation setup. The experimental unit is a single animal.

### 2.2 Lewis Lung Carcinoma

Lewis Lung Carcinoma (LLC) cancer was induced as previously described ^3^. LLC cells (ATCC® CRL1642™) were cultured in DMEM, high glucose (Gibco, #41966-029, USA) supplemented with 10% fetal bovine serum (FBS, Sigma-Aldrich, #F0804, USA), 1% penicillin-streptomycin (ThermoFisher Scientific, #15140122, USA) (5% CO2, 37 °C). Prior to inoculation into mice, LLC cells were trypsinized and washed twice with PBS. LLC cells were suspended in PBS with a final concentration of 2.5 * 10^6^ cells/ml. All mice were shaved on the flank two days prior to the inoculation and randomized into two groups with similar average body weight. The mice were subcutaneously injected with PBS with or without 2.5 * 10^5^ LLC cells into the right flank. The experiments were carried out 19 and 21 days after cancer cell inoculation. Mice developing ulcerations (human endpoint) were sacrificed by cervical dislocation. Mice with tumor >0.5 gram were excluded. Three animals were excluded due to the size of the tumor.

### 2.3 Ex vivo muscle incubations

On the day of experimentation, fed mice were anaesthetized by intraperitoneal injection of pentobartital/lidocain (6 mg of pentobarbital sodium and 0.6 mg of lidocain/100 g of body weight) after which soleus and EDL muscles were tied with non-absorbable 4-0 silk suture loops (Look SP116, Surgical Specialities Corporation) at both ends and suspended between adjustable hooks at resting length (1-2 mN tension) in ex vivo incubation chambers (Multi Myograph system, Danish Myo-Technology) at 30°C with continuously 95% O2/5% CO2-bubbled Krebs-Ringer-Henseleit (KRH) buffer (118.5 mM NaCl, 24.7 mM NaHCO3, 4.74 mM KCl, 1.18 mM MgSO4·7H2O, 1.18 mM KH2PO4, 2.5 mM CaCl2·2H2O) supplemented with 8 mM mannitol and 2 mM pyruvate (KRH medium). The experimental groups were randomized between chambers. The tumors and spleens were also dissected at this stage, rinsed and snap frozen in liquid nitrogen, before the mice were sacrificed by cervical dislocation. After dissection, the muscles were first allowed 15 min of recovery in fresh KRH buffer and then incubated for 10 min in KRH with or without 1.5 nM of insulin (sub-maximal dose). Next, the medium was changed to one containing radioactively labelled 2-[3H] deoxyglucose (2-DG; 0.30 μCi/ml in 1 mM non-radiolabelled 2-DG) and mannitol (0.28 μCi/ml in 8 mM non-radiolabelled mannitol) and 10 min of tracer labelling were allowed. For the insulin-stimulated group, the same insulin concentration was maintained in the tracer medium. Finally, the muscles were harvested, rinsed in ice-cold KRH medium, dabbed dry on paper and snap-frozen in liquid nitrogen until further analysis.

### 2.4 Immunoblotting and glucose transport measurements

The frozen soleus and EDL muscles were trimmed free of connective tissue and sutures and weighed. Muscles were homogenized 1 min at 30 Hz using a TissueLyser II bead mill (Qiagen, USA) in 300 μl ice-cold homogenization buffer, pH 7.5 (10% glycerol, 1% NP-40, 20 mM sodium pyrophosphate, 150 mM NaCl, 50 mM HEPES (pH 7.5), 20 mM β-glycerophosphate, 10 mM NaF, 2 mM phenylmethylsulfonyl fluoride, 1 mM EDTA (pH 8.0), 1 mM EGTA (pH 8.0), 2 mM Na3VO4, 10 μg/ml leupeptin, 10 μg/ml aprotinin, 3 mM benzamidine). After the homogenization, the samples were rotated end-over-end for 30 min at 4 °C, before being subjected to centrifugation (9500 RCF) for 20 min at 4 °C. The lysates were then collected. Lysate protein concentrations were measured using the bicinchoninic acid method with bovine serum albumin (BSA) as a standard. A fraction (50 μl) of the lysate was dissolved in 2 ml of β-scintillation liquid (Ultima Gold, Perkin Elmer) for measurement of 2-DG transport using [14C]mannitol to estimate extracellular space using β-scintillation counting. The 2-DG transport was related to the protein concentration of the lysate. The measurements of 2-DG transport were blinded.

The remaining lysate were used for standard immunoblotting of total proteins and phosphorylation levels of relevant proteins. In brief, Polyvinylidene difluoride membranes (Immobilon Transfer Membrane; Millipore) were blocked in Tris-buffered saline (TBS)-Tween 20 containing 2% skim milk or 3% bovine serum albumin (BSA) for 5 min at room temperature. Membranes were incubated with primary antibodies (Table 1) overnight at 4 °C, followed by incubation with HRP-conjugated secondary antibody for 45 min at room temperature.

**Table 1:** Antibodies.

| Protein | Company | Catalog # | Dilution |
| --- | --- | --- | --- |
| Akt2 | Cell signaling technology | 3063 | 1:1000, 2% Skim milk |
| pAkt ser473 | Cell signaling technology | 9271 | 1:1000, 2% Skim milk |
| pAkt thr308 | Cell signaling technology | 9275 | 1:1000, 2% Skim milk |
| TBC1D4 | Abcam | ab189890 | 1:1000, 2% Skim milk |
| pTBC1D4 thr642 | Cell signaling technology | 8881 | 1:1000, 2% Skim milk |
| pTBC1D4 ser588 | Cell signaling technology | 8730 | 1:1000, 2% Skim milk |
| pTBC1D4 ser318 | Cell signaling technology | 8619 | 1:1000, 2% Skim milk |
| p70S6K total | Cell signaling technology | 2708 | 1:1000, 2% Skim milk |
| p p70S6K thr389 | Cell signaling technology | 9205 | 1:1000, 2% Skim milk |
| Ribosomal protein S6 (rS6) | Cell signaling technology | 2217 | 1:1000, 3% Skim milk |
| rS6 ser235/ser236 | Cell signaling technology | 2211 | 1:1000, 3% BSA |
| Hexokinase II | Cell signaling technology | 2867 | 1:1000, 2% Skim milk |
| GLUT4 | Thermo Fisher Scientific | PA1-1065 | 1:1000, 2% Skim milk |
| Pyruvate dehydrogenase | D.G. Hardie (University of Dundee, Scotland) | - | 1 µg/mL, 2% Skim milk |
| AMPK alpha2 | Abcam | 3760 | 1:1000, 2% Skim milk |
| pAMPK thr172 | Cell signaling technology | 2531 | 1:1000, 2% Skim milk |
| ACC total | Dako – streptavidin | P0397 | 1:2500, 3% BSA |
| pACC ser212 | Cell signaling technology | 3661 | 1:1000, 2% Skim milk |
| ULK1 | Cell signaling technology | 8054 | 1:500, 3% BSA |
| pULK ser757 | Cell signaling technology | 6888 | 1:1000, 2% Skim milk |
| mTOR total | Cell signaling technology | 2983 | 1:1000 2 % skim milk |
| p-mTOR ser2448 | Cell signaling technology | 2971 | 1:1000 2 % skim milk |
| Rodent Oxphos | Abcam | ab110413 | 1:5000, 2% Skim milk |
| FATP4 | Abcam | ab200353 | 0.442µg/µL, 2% Skim milk |
| Citrate Synthase | Abcam | ab96600 | 1:5000, 3% BSA |
| Glycogen Synthase | Gift from Prof. Oluf Pedersen | - | 1:20000 2 % skim milk |
| GSK3 beta | BD Bioscience | 610202 | 1:1000 2 % skim milk |
| pGSK3 alpha/beta ser21/9 | Cell signaling technology | 9331 | 1:1000, 2% Skim milk |

Coomassie brilliant blue staining was used as a loading control^17^. The same coomassie brilliant blue staining is presented, when the same four samples are presented for several proteins investigated using the same membrane (e.g. Fig. 3). To ensure quantification within the linear range for each antibody probed, a standard curves were made for total proteins, basal and insulin-stimulated conditions. Bands were visualized using the Bio-Rad ChemiDoc MP Imaging System and enhanced chemiluminescence (ECL+; Amersham Biosciences). Bands were quantified using Bio-Rad’s Image Lab software 6.0.1.

### 2.5 Statistics

All statistics were performed using GraphPad Prism, 8.0 (GraphPad Software, La Jolla, CA, USA). Statistical testing was performed using student’s t-test and two-way repeated measures ANOVA (the two EDL muscles and the two soleus muscles from the same mouse were treated as pairs comparing basal vs. insulin stimulation) as applicable. The main effects and interactions are presented in the figures when significant. For post-hoc analyses, a Sidak’s multiple comparisons test was performed. The significance level was set at α<0.05.

### 2.6 Data presentation and graphics

All graphs were created using GraphPad Prism, 9.0 (GraphPad Software, La Jolla, CA, USA). All figures were created using Inkscape (Inkscape.org). Illustrations were created using ©BioRender.com.

## 3.0 Results

### 3.1 Cancer leads to a minor reduction in GLUT4 protein content in soleus muscle

At day 19-21 post tumor inoculation, soleus and EDL muscles were isolated, incubated, and stimulated with or without a submaximal concentration of insulin (Fig. 1A). On the experimental day, the average tumor size was ~2.0 gram (Fig. 1B) and body mass tended lower (p=0.0867) in tumor-bearing mice (Fig. 1C). Spleen weight was increased (+145%, Fig. 1D), indicative of pre-cachexia and elevated inflammation in tumor-bearing mice compared to controls.

**Figure 1:**
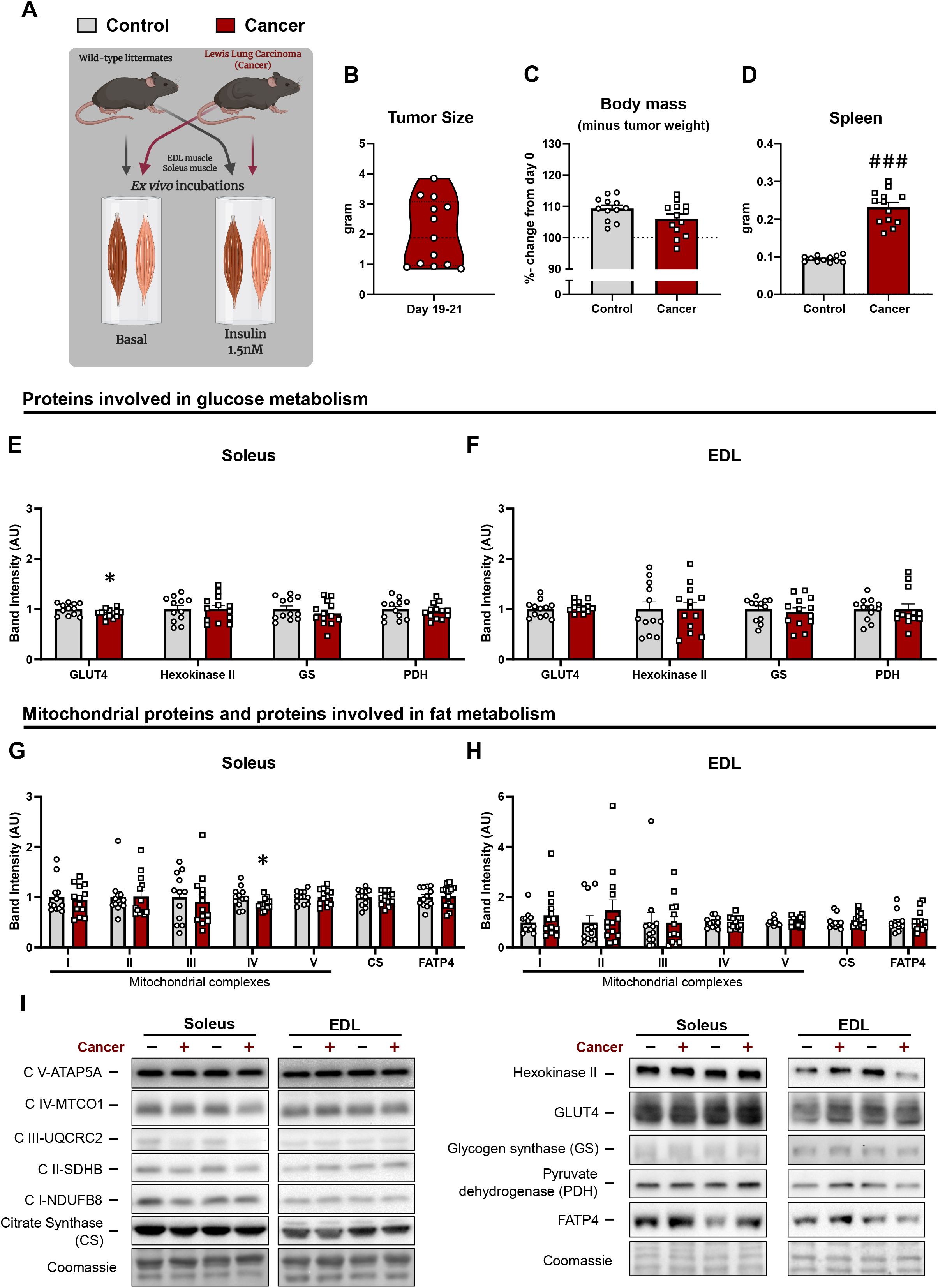
The effect of Lewis Lung Carcinoma cells inoculation on tumor and spleen weight, body mass and protein expression of proteins involved in glucose and fat metabolism. A) C57BL/6J mice were subcutaneously inoculated with Lewis Lung Carcinoma (LLC) cells (cancer) or saline (control) along the flank. Nineteen-21 days later the mice were sacrificed and soleus and extensor digitorum longus (EDL) muscles were incubated ex vivo. B) Tumor weight in LLC inoculated mice. C) Change (%) in body mass from the day of tumor inoculation (day 0). D) Spleen mass. Muscle protein expression of proteins involved in glucose metabolism in E) soleus and F) EDL, as well as mitochondrial proteins and proteins involved in fat metabolism in G) soleus, and H) EDL. I) Representative western blots of investigated proteins. For control mice; n=12, for tumor-bearing (cancer) mice; n=13. Effect of cancer; # / ### = p<0.05 / p<0.001.

We firstly investigated key proteins related to glucose transport and mitochondrial proteins, namely glucose transporter 4 (GLUT4), hexokinase II (HK II), glycogen synthase (GS), pyruvate dehydrogenase (PDH), subunits of the electron transport chain (ETC), citrate synthase (CS), and long-chain fatty acid transport protein 4 (FATP4). Cancer lead to a minor reduction in protein content of GLUT4 (−9%) and complex 4 of the ETC (−13%) in soleus muscle of tumor-bearing mice compared to control mice. No effects of cancer were observed on the other proteins investigated in either muscles (GLUT4; Fig. 1E/F, HK II; Fig. 1E/F, GS; Fig. 1E/F, PDH; Fig. 1E/F, ETC; Fig. 1G/H, CS; Fig. 1G/H and FATP4; Fig. 1G/H). Representative western blots are shown in Fig. 1I. Collectively, no major changes were observed for key proteins related to glucose handling.

### 3.2 Cancer selectively causes insulin resistance in oxidative soleus muscle

We next investigated the glucose transport during submaximal (1.5 nM) insulin stimulation. As expected, insulin increased glucose transport in both soleus (+115%, Fig. 2A/B) and EDL (+55%, Fig. 2C/D) muscles from non-tumor-bearing control mice. Remarkably, this response was abrogated in the oxidative soleus muscle of tumor-bearing mice (Fig. 2A/B). This effect was muscle-type specific, as insulin increased glucose transport by 70% in the glycolytic EDL muscle from tumor-bearing mice with no effect of cancer (Fig. 2C/D). These data demonstrate that cancer affects muscles differently dependent on muscle-type; oxidative or glycolytic. We subsequently investigated insulin signaling pathways (Fig. 2E), in order to determine the molecular underpinnings of the different response to cancer in muscle.

**Figure 2:**
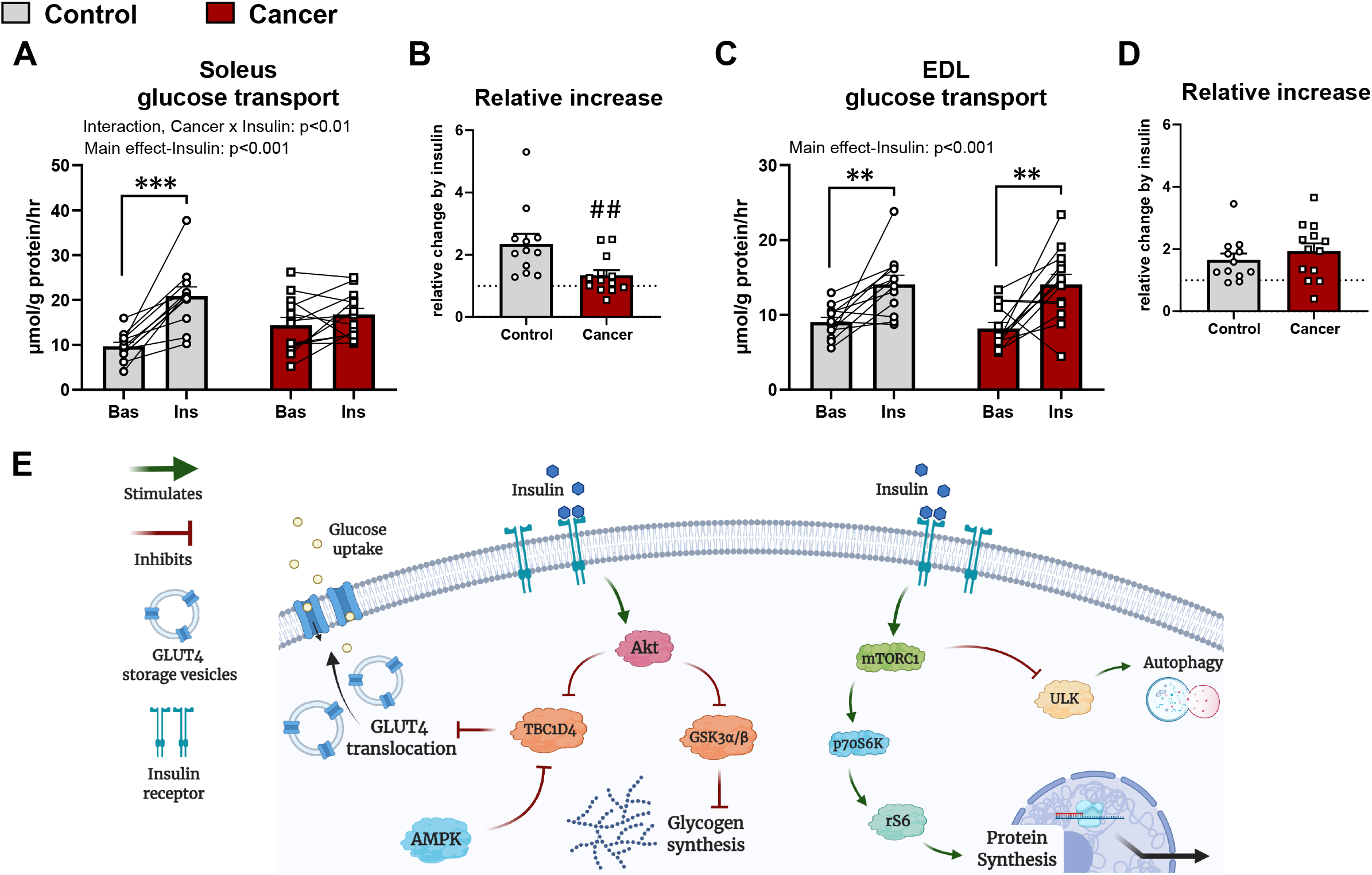
Muscle glucose transport is affected in the oxidative soleus muscle of tumor-bearing mice, not the glycolytic extensor digitorum longus muscle. A) 2-deoxyglucose transport in soleus muscles stimulated with or without insulin (1.5 nM), and B) the relative effect of insulin on glucose transport. C) 2-deoxyglucose transport in extensor digitorum longus (EDL) muscles stimulated with or without insulin (1.5 nM), and D) the relative effect of insulin on glucose transport. E) Schematic illustration of the insulin signaling pathways investigated in current study. For control mice; n=12, for tumor-bearing mice; n=13. The connecting lines illustrate muscles from the same mouse (Basal (Bas) vs. insulin (Ins)). Effect of insulin; **/ *** = p<0.01 / p<0.001. Effect of cancer; ## = p<0.01.

### 3.3 Cancer inhibits insulin-stimulated TBC1D4 and GSK3 phosphorylation

Proximal insulin signaling via phosphorylation (p) of Akt threonine(thr)308 (Fig. 3A) and pAkt serine(ser)473 (Fig. 3B)was similarly increased by insulin in control and tumor-bearing mice, . The Rab GTPase activating protein TBC1D4, downstream of Akt, is inactivated by phosphorylation, which is necessary for translocation of GLUT4 to the plasma membrane^18^. As TBC1D4 has multiple insulin-sensitive phosphorylation sites, we measured if any alteration in these phospho-sites could explain the lack of effect of insulin on glucose uptake in soleus muscle. More specifically, we investigated pTBC1D4 ser318, ser588, and thr642 (in mice; ser324, ser595, thr649), which are all phosphorylated during insulin stimulation^19,20^, and are direct targets of Akt, but also other kinases^21^. In soleus muscle of control mice, insulin led to a 35% increased phosphorylation of ser318 (Fig. 3C), 60% increased ser588 (Fig. 3D), and 150% increased thr642 (Fig. 3E) of TBC1D4. In contrast, none of these phosphorylations were increased during insulin stimulation in soleus muscle from tumor-bearing mice (Fig. 3C, 3D, and 3E). In addition, basal pTBC1D4 at ser588 (Fig. 3D) and thr642 (p=0.086, Fig. 3E) were increased or tended to be increased, respectively, in soleus of tumor-bearing mice compared to control mice. This was in contrast to EDL, where insulin increased the phosphorylation of all the above mentioned phospho-sites independent of cancer (Fig. 3C, 3D, and 3E). In fact, the post-hoc test demonstrated that the insulin-effect on TBC1D4 ser588 and thr642 was driven by the increase in the EDL muscles from the tumor-bearing mice.

**Figure 3:**
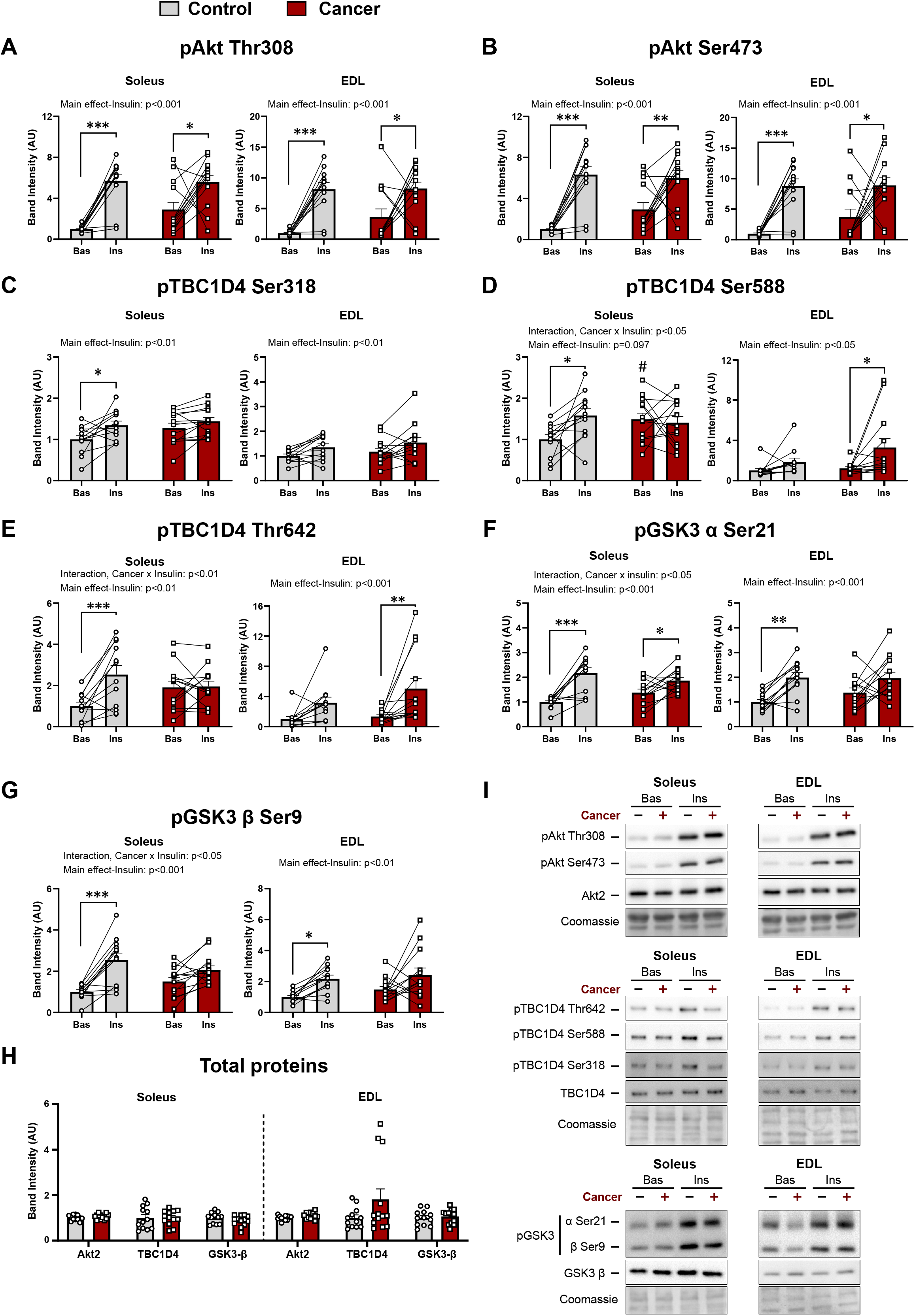
Cancer abrogates the insulin-stimulated phosphorylation of TBC1D4 in soleus muscle. Quantification of A) phosphorylated (p) Akt thr308, B) pAkt ser473, C) pTBC1D4 ser318, D) pTBC1D4 ser588, E) pTBC1D4 thr642, F) pGSK3-α ser21, G) pGSK3-β ser9, and H) total proteins of Akt2, TBC1D4 and GSK3-β. I) Representative western blots of investigated proteins. For control mice; n=12, for tumor-bearing (cancer) mice; n=13. The connecting lines illustrate muscles from the same mouse (Basal (Bas) vs. insulin (Ins)). Effect of insulin; * / ** / *** = p<0.05 / p<0.01 / p<0.001. Effect of cancer; # = p<0.05.

Thus, these data show that in soleus muscle, cancer impairs insulin signal transduction to TBC1D4 on several phosphorylation sites, which could explain the reduced insulin-stimulated glucose uptake observed in soleus muscle of tumor-bearing mice. In contrast, no impairment of TBC1D4 phosphorylation was observed in EDL muscles from tumor-bearing mice that did not display any alterations in glucose uptake compared to control mice.

To test whether this phenotype transferred to other Akt substrates, we investigated the phosphorylation of glycogen synthase kinase 3 (GSK3) α/β^22,23^. GSK3α/β negatively regulate the protein glycogen synthase (synthesis of glycogen). Thus, phosphorylations of GSK3α/β (ser21/ser9) inhibit the kinase activity of glycogen synthase and thereby promote glycogen synthesis^24,25^. In control mice, insulin increased phosphorylation of GSK3 α/β in both soleus (α: 115% and β: 155%, Fig. 3F/G) and EDL muscle (α: 100% and β: 120%, Fig. 3F/G). In both soleus from tumor-bearing mice, insulin-stimulated GSK3 α/β appeared diminished. Here, GSK3 α phosphorylation increased by 35% in response to insulin (Fig. 3F), and GSK3 β phosphorylation tended (p=0.097) to increase (Fig. 3G). In EDL muscle from tumor-bearing mice, GSK3 phosphorylation-sites (α: 40%, p=0.07, and β: 65%, p=0.072) tended increase in response to insulin (Fig. 3F/G). Total protein content of Akt2, TBC1D4, and GSK3 β protein content were similar between control and tumor-bearing mice in both soleus and EDL muscle (Fig. 3H). Thus, cancer impaired GSK3 phosphorylation in both soleus muscle, indicative of defective glycogen synthesis in both insulin resistant soleus muscle and insulin sensitive EDL muscle. Representative western blots are shown in fig. 3I.

### 3.4 Cancer promotes AMPK activation in EDL, but not soleus, muscle

AMP-activated protein kinase (AMPK) is metabolic stress-sensor in muscle^26^ that is proposed to be involved in glucose uptake in response to exercise^27–30^. AMPK also provide input to insulin signaling and is required for the increase in insulin-sensitivity after muscle contraction^31,32^. AMPK phosphorylates TBC1D4 on ser588^21^, which was upregulated in EDL muscles from tumor-bearing mice (Fig. 3D) and we therefore investigated AMPK signaling.

pAMPK thr172 (Fig. 4A) and pACC ser212 (a direct AMPK substrate) (Fig. 4B) were similar between control and tumor-bearing mice in soleus muscle. In contrast, both AMPK and ACC phosphorylations were upregulated in the EDL muscle of tumor-bearing mice (Fig. 4A and 4B), aligning with previous reports^33,34^. Total AMPK α2 and ACC1/2 (Fig. 4C) protein content were not affected by cancer. Representative western blots are shown in fig. 4D. Thus, elevated AMPK activation might be involved in the protection from cancer-induced insulin resistance in EDL muscle.

**Figure 4:**
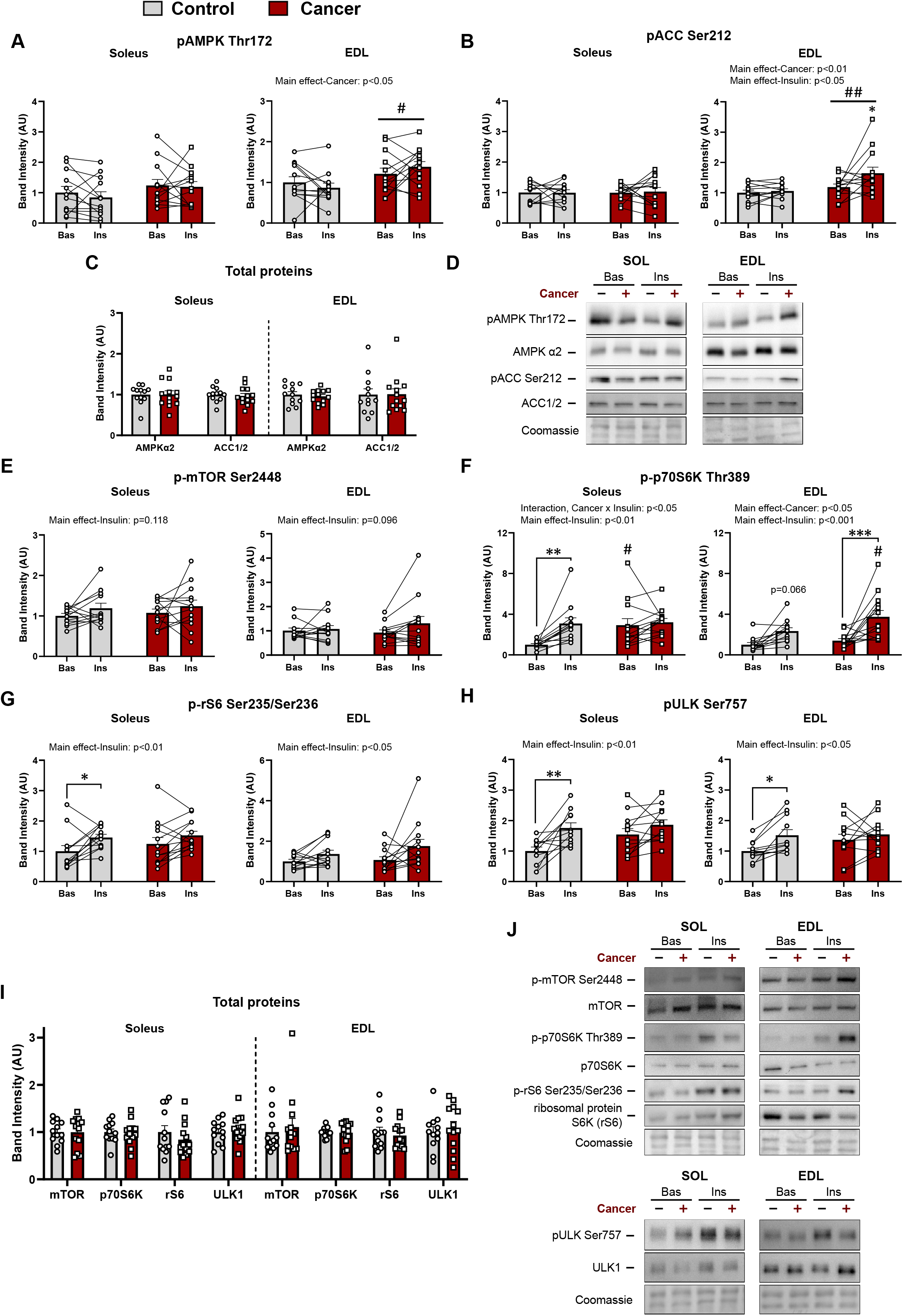
Cancer leads to disrupted or reduced mTORC1 signaling during insulin stimulation. Quantification of A) phosphorylated (p) AMPK thr172, B) pACC ser212, and C) total AMPK α2 and total ACC1/2 protein expression in both soleus and extensor digitorum longus (EDL) muscles. D) Representative western blots of phosphorylated and total AMPK and ACC. mTORC1 signaling was measured via phosphorylation of E) mTOR ser2448, F) p-p70S6K thr389, G) p-rS6 ser235, and H) pULK ser757. I) Total proteins of mTOR, p70S6K, rS6, ULK1. J) Representative western blots of investigated proteins in E-I. For control mice; n=12, for tumor-bearing mice; n=13, except for H) soleus (control mice; n=11, for tumor-bearing mice; n=12). The connecting lines illustrate muscles from the same mouse (Basal (Bas) vs. insulin (Ins)). Effect of insulin; * / **/ *** = p<0.05 / p<0.01 / p<0.001. Effect of cancer; # / ## = p<0.05 / p<0.01.

### 3.5 Cancer altered mTORC1 signaling in both soleus and EDL muscle

Mammalian target of rapamycin complex 1 (mTORC1) is a central regulator of cell size and protein synthesis^35^. Cancer can lead to decreased protein synthesis in human muscle^36–38^, and reduced/abrogated mTORC1 signaling has been observed in various pre-clinical cancer mouse models^33,34,39–44^. Thus, we next determined the effect of cancer on insulin-stimulated anabolic signaling in muscle.

Insulin-stimulated phosphorylation of mTOR at ser2448, a reported insulin-sensitive site^45^, was not affected by either sub-maximal insulin or cancer in soleus and EDL muscle (Fig. 4E). Despite no increase in mTOR phosphorylation, insulin increased phosphorylation of the downstream target of mTORC1, p70S6K thr389, in both soleus (+208%) and EDL (+134%, p=0.066) of control animals (Fig. 4F). This effect of insulin on p-p70S6K thr389 was completely abrogated in soleus muscle of tumor-bearing mice compared to control mice (Fig. 4F). In contrast, p-p70S6K thr389 was augmented in tumor-bearing mice during insulin stimulation compared to control mice in EDL muscle (+60%, Fig. 4F). p70S6K activity leads to phosphorylation of ribosomal protein S6 (rS6) at ser235/236^46^. In soleus muscle, insulin only led to an increase in phosphorylation in control animals, not tumor-bearing mice (+45%, Fig. 4G) as seen for p-p70S6K thr389. In EDL muscle, insulin caused a main effect of increased p-rS6 ser235 with no effect of cancer (Fig. 4G). ULK1 is another downstream target of mTORC1 and phosphorylation of ULK1 at ser757 leads to inhibition of autophagy^47^ (Fig. 2E). Interestingly, phosphorylation of ULK1 at ser757 was abrogated in both soleus and EDL muscle of tumor-bearing mice, where this site increased in both muscles during insulin stimulation in control mice (soleus: +75%, EDL: +52%) (Fig. 4H). Thus, insulin leads to the phosphorylation of ULK, but seemingly not in tumor-bearing mice. Total mTOR, p70S6K, rS6, and ULK1 (Fig. 4I) protein content were unaffected by cancer. Representative western blots are shown in Fig. 4J.

Taken together, these results suggest that the observed cancer-induced impairment of glucose transport in soleus muscle also manifested in anabolic resistance indicated by disrupted mTORC1 downstream signaling during insulin stimulation.

## 4.0 Discussion

Here, we present evidence of selective insulin resistance within different muscle-types in response to cancer in mice (Fig. 5). A primary finding was that cancer prevented insulin-stimulated glucose transport in oxidative soleus muscle, but not glycolytic EDL muscle. Secondly, this selective insulin resistance was associated with an inability for insulin to elicit multi-site phosphorylation of the Rab GTPase activating protein, TBC1D4 and GSK3, despite full induction of pAkt. Thirdly, we found that insulin-stimulated mTORC1 signaling, p70S6K-S6-and ULK-signaling, were abrogated by cancer. Collectively, these data show that cancer selectively rewires oxidative soleus muscle causing severe insulin resistance, which could lead to the metabolic dysregulation observed in cancer.

**Figure 5:**
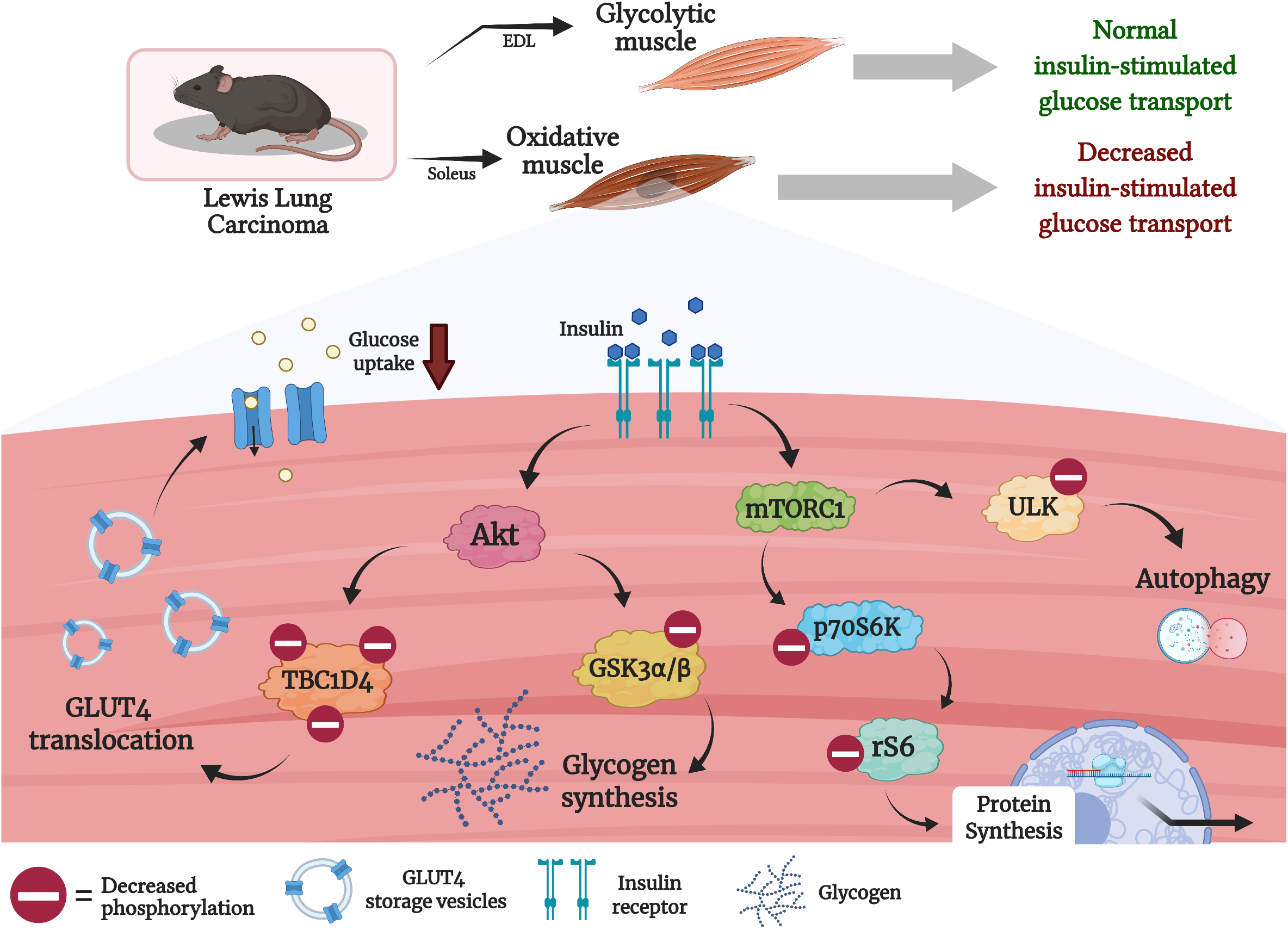
Schematic illustration of the data presented in current study.

Our discovery that cancer abrogates insulin-stimulated glucose transport in soleus muscle expands on other studies that have reported reduced blood glucose-lowering effect of insulin in vivo of tumor-bearing rodents^2–4^ and in patients with cancer^5–9,48^. Such findings are clinically relevant, because metabolic disturbances are associated with cancer incidence, poor cancer prognosis, and increased recurrence rates^10–15^. Whole body insulin resistance, measured by hyperinsulinemic-euglycemic clamp has been reported in cancers such as gastrointestinal^6,8,48^, colorectal^5,8,48^, lung^8,9,48^, and pancreatic cancer^7^. Based on our present results as well as a recent study^3^, whole body insulin resistance and glucose intolerance in many cancers are likely due to abrogated skeletal muscle glucose uptake. The results of the current investigation would suggest that distorted insulin signaling in muscle leads to insulin resistance specifically in oxidative muscles. In agreement with this observation, proteomic analyses of human^49^ and rodent^50,51^ skeletal muscle show that proteins involved in oxidative metabolism are highly altered in cancer cachexia.

A second important finding was that cancer-associated insulin resistance in soleus was accompanied by dysregulation on multiple phosphorylation-sites on TBC1D4 (ser318, ser588, and thr642), of which thr642 previously has been shown to be important for insulin-stimulated glucose uptake in skeletal muscle^18,52^. Interestingly, tumor-bearing mice displayed normal phosphorylation of Akt, which phosphorylates TBC1D4 at thr642. Thus, we speculate that the signal transduction from Akt to TBC1D4, TBC1D4 itself or TBC1D4 phosphatases are dysregulated in oxidative muscle of tumor-bearing mice. Similarly to our findings, phosphorylation at several sites on TBC1D4, including ser318, ser588, and ser751, are impaired during insulin stimulation in muscles from patients with T2D^53^, suggesting that reduced TBC1D4 signaling can be a common trait in insulin resistant skeletal muscle. Likewise, T2D has been associated with mild reductions in skeletal muscle expression of GLUT4 protein^54,55^, which was also observed in soleus muscle of tumor-bearing mice, but this is not always observed in T2D^53^. Reduced GLUT4 protein content align with TBC1D4 dysregulation as lack of TBC1D4 or loss-of-function mutants result in reduced GLUT4 content in mouse skeletal muscle^56^ and human muscle^57^.

In contrast to the insulin resistant soleus muscle of tumor-bearing mice, the insulin sensitive EDL muscle displayed normal insulin-induced TBC1D4 phosphorylation and cancer had no effects on GLUT4 protein content. In fact, TBC1D4 ser588 was elevated in EDL muscles of tumor-bearing mice. AMPK is a kinase for TBC1D4 including at the ser588 site^21^, and we speculate that the elevated AMPK signaling in EDL muscles from tumor-bearing mice may be a compensatory mechanism that preserves insulin sensitivity. Notably, AMPK is a positive regulator of insulin sensitivity via TBC1D4 after muscle contractions^31,32^ and AMPK seems to be required for normal insulin-induced signaling of the TBC1D4 paralogue and Rab GTPase activating protein, TBC1D1, in mouse muscle^58^. Our findings thus identify an intriguing link between AMPK and insulin sensitivity in the context of cancer that should be explored in future studies.

A third major finding was that insulin-stimulated p-p70S6K thr389, p-rS6 ser235/236, and pULK ser757 were abolished in soleus muscle of tumor-bearing mice, indicative of reduced mTORC1 signaling and anabolic resistance. mTORC1 signaling in skeletal muscle has previously been reported to be decreased in cancer cachexia at baseline^33,39–41^, during contraction^40,42^, and after an intraperitoneal glucose injection^34^. Yet, other studies show unchanged or increased mTORC1 signaling in cachectic rodent models^59^ and humans^60^. Our study show that altered mTORC1 activity also extends to insulin-stimulated mTORC1 signaling and suggests that cancer-associated insulin resistance extends to the level of anabolism. The current data support the theory that cancer leads to muscle insulin resistance^61–63^, which in turn could accelerate muscle loss in cancer cachexia. However, this has yet to be experimentally verified in pre-clinical models or patients.

In conclusion, we show that cancer leads to marked insulin resistance in oxidative mouse soleus muscle evidenced by blocked insulin-stimulated glucose transport and abolished insulin-induced phosphorylation of TBC1D4 and GSK3 at multiple phosphorylation sites. Furthermore, cancer impaired mTORC1 signaling, measured via p70S6K-rS6 and ULK1 phosphorylation, in soleus muscle, while only ULK1 phosphorylation was impaired in EDL muscle of tumor-bearing mice. Our results shows how cancer leads to insulin resistance in a muscle-type specific manner, and we identify the potential molecular mechanisms leading to this phenotype, which could guide future studies and optimize cancer therapy.

## CRediT authorship contribution statement

**Steffen H. Raun**: Conceptualization, Methodology, Validation, Formal analysis, Investigation, Writing - Original Draft, Visualization, Project administration. **Jonas Roland Knudsen**: Conceptualization, Methodology, Validation, Formal analysis, Investigation, Writing - Original Draft. **Thomas E. Jensen**: Investigation, Writing - Review & Editing. **Xiuqing Han**: Investigation, Writing - Review & Editing. **Lykke Sylow**: Conceptualization, Methodology, Writing - Original Draft, Supervision, Project administration, Funding acquisition.

## Acknowledgement

We would like to acknowledge Nicoline Resen Andersen for the help with running parts of the immunoblotting in current study.

## Data access

For question(s) or access to data, please contact corresponding author Lykke Sylow.

